# Effects of MP2RAGE B_1_^+^ sensitivity on inter-site T_1_ reproducibility and morphometry at 7T

**DOI:** 10.1101/2020.02.13.947382

**Authors:** Roy AM Haast, Jonathan C Lau, Dimo Ivanov, Ravi S Menon, Kâmil Uludağ, Ali R Khan

## Abstract

Most neuroanatomical studies are based on MR images, whose intensity profiles are not solely determined by the tissue’s longitudinal relaxation times (T_1_) but also affected by varying non-T_1_ contributions, hampering data reproducibility. In contrast, quantitative imaging using the MP2RAGE sequence, for example, allows direct characterization of the brain based on the tissue property of interest. Combined with 7 Tesla (7T) MRI, this offers unique opportunities to obtain robust high-resolution brain data characterized by a high reproducibility, sensitivity and specificity. However, specific MP2RAGE parameters choices – e.g., to emphasize intracortical myelin-dependent contrast variations – can substantially impact image quality and cortical analyses through remnants of B_1_^+^-related intensity variations, as illustrated in our previous work. To follow up on this: we (1) validate this protocol effect using a dataset acquired with a particularly B_1_^+^ insensitive set of MP2RAGE parameters combined with parallel transmission excitation; and (2) extend our analyses to evaluate the effects on hippocampal and subcortical morphometry. The latter remained unexplored initially but will provide important insights related to generalizability and reproducibility of neurodegenerative research using 7T MRI. We confirm that B_1_^+^ inhomogeneities have a considerably variable effect on cortical T_1_ and thickness estimates, as well as on hippocampal and subcortical morphometry depending on MP2RAGE setup. While T_1_ differed substantially across datasets initially, we show inter-site T_1_ comparability improves after correcting for the spatially varying B_1_^+^ field using a separately acquired Sa2RAGE B_1_^+^ map. Finally, as for cortical thickness, removal of B_1_^+^ residuals affects hippocampal and subcortical volumetry and boundary definitions, particularly near structures characterized by strong intensity changes (e.g. cerebral spinal fluid and arteries). Taken together, we show that the choice of MP2RAGE parameters can impact T_1_ comparability across sites and present evidence that hippocampal and subcortical segmentation results are modulated by B_1_^+^ inhomogeneities. This calls for careful (1) consideration of sequence parameters when setting acquisition protocols; as well as (2) interpretation of results focused on neuroanatomical changes due to disease.

**Highlights:**

- Previously observed effects of B_1_^+^ inhomogeneities on cortical T_1_ and thickness depend strongly on MP2RAGE parameters
- Inter-site comparability of cortical T_1_ and thickness greatly improves after removal of B_1_^+^ residuals
- Post-hoc MP2RAGE B_1_^+^ correction affects hippocampal (and subcortical) size and shape analyses
- Neuroradiological research would benefit from careful examination of imaging protocols and their impact on results, especially when B_1_^+^ maps are not acquired

## 1. Introduction

Magnetic resonance imaging (MRI) at 7 Tesla (7T) and its established increase in sensitivity and specificity allows characterization of the brain with a level of detail that cannot readily be obtained at lower field strengths (Uğurbil, 2018). But despite its promises, several data quality issues, limiting data interpretation and reproducibility, are still hindering complete acceptance of 7T MRI into clinical practice. Quality assessment, standardization and harmonization of 7T MRI protocols are increasingly becoming appreciated by the neuroimaging field to allow utilization and generalization of protocols across studies, imaging sites and scanner vendors (Poldrack et al., 2017). Several nationwide (Clarke et al., 2019; Voelker et al., 2016) and international (Düzel et al., 2019) initiatives have embarked on such establishments indicating the importance of this issue. These consortia aim to set up standardized sequences across the main 7T MRI vendors to limit the long-known effects of hard- (e.g. coils and gradients) and software (e.g. imaging sequence implementations and reconstruction methods) differences on MRI analyses (Jovicich et al., 2009). As such, large population imaging studies, too expensive to cover by individual institutions, as well as those focusing on rare diseases, will benefit by allowing data pooling across multiple imaging sites.

In essence, to improve reproducibility, sequences and corresponding parameters need to be chosen in such a way that they provide robust data characterized by comparable temporal and spatial signal-to-noise (SNR) and contrast-to-noise (CNR) ratios, as well as intensity profiles, independent of acquisition site, scanner vendors and/or time point (Voelker et al., 2016). Quantitative MRI (ideally) overcomes potential, non-biochemical inter-site, intra-subject biases that are present in weighted MRI data (Haast et al., 2016; Okubo et al., 2016; Weiskopf et al., 2013), which hinder the direct comparison across studies and between patients and healthy controls. There are numerous options for quantitative imaging in terms of sequences, depending on the tissue property of interest. The MP2RAGE (magnetization-prepared two rapid gradient echo) sequence gained significant popularity during the last decade (Marques et al., 2010). It is widely used for anatomical imaging as it provides a ‘standard’ T_1_-weighted (T_1_w) image and allows quantification of the longitudinal relaxation time (T_1_), ideally free of T_2_*, M_0_ (i.e., proton density) and B_1_^-^ effects. These images support analyses using conventional analysis tools, such as FreeSurfer (Fischl, 2012) or FSL-FIRST (Patenaude et al., 2011), for assessment of the brain’s morphology, along with T_1_ relaxometry to study biochemical (mostly myelin-dependent, Stüber et al. (2014)) changes due to learning, aging and/or disease.

While the MP2RAGE approach eliminates potential biases present in non-quantitative imaging methods, residual biases related to the radio frequency transmit field (B_1_^+^) may persist. Importantly, when setting up an imaging protocol, MP2RAGE parameters can be set to render images minimally sensitive to spatial variations in transmit efficiency (i.e., B_1_^+^ field) or to sensitize images for (i.e., myelination-dependent) contrast variations within subcortical and/or cortical tissue while risking B_1_^+^ residues. These transmit efficiency (B_1_^+^) variations – too strong to correct for using dielectric pads (Teeuwisse et al., 2012) – introduce image artifacts that hamper accurate T_1_ estimation and subsequent analyses in the affected regions. For example, we have established earlier that accuracy of automatic neocortical segmentation using FreeSurfer based on 7T MP2RAGE data suffers from severe local B_1_^+^ field inhomogeneity effects near inferior temporal and frontal lobes (Haast et al., 2018b). As a result, tedious manual corrections of the automatic image segmentations would be necessary to correct for these tissue classification errors. Post-hoc removal of the residual B_1_^+^ inhomogeneities using a separately acquired B_1_^+^ map (Eggenschwiler et al., 2012; Marques and Gruetter, 2013), showed to be capable to reduce the cortical T_1_ and thickness quantification errors substantially. However, it remained unclear how the corrected average cortical T_1_ and thickness estimates directly compared against an independent dataset obtained within a setting that would provide MP2RAGE images already minimally sensitive to B_1_^+^ variations. In addition, analyses were restricted to cortical gray matter and did not characterize the effects on the delineation of hippocampal and subcortical gray matter.

As such, we first aim to replicate our initial neocortical analyses using an independent MP2RAGE dataset acquired at 7T at the Centre for Functional and Metabolic Mapping (CFMM) in London (Ontario, Canada) to highlight the effect of MP2RAGE B_1_^+^ sensitivity on the previously observed changes in cortical T_1_ and thickness (i.e., inter-site comparison). We used sequence parameters more closely matching those proposed in Marques and Gruetter (2013), lowering B_1_^+^-sensitivity, and used a parallel transmit head coil to increase B_1_^+^ homogeneity. Secondly, using morphometric analyses we characterize the effect of the residual B_1_^+^ field on FreeSurfer’s hippocampal and subcortical segmentations. While our analyses will cover the thalamus, caudate nucleus, putamen, globus pallidus and nucleus accumbens, we focus on hippocampal morphometry in particular, and analyse volume and shape differences between original and the post-hoc corrected data. The hippocampus is one of the most studied structures of the brain and plays an important role in the functioning of the brain’s learning and memory system (Small et al., 2011). Therefore, insights into the magnitude of volume and shape differences induced by B_1_^+^-inhomogeneities may have important implications for the reproducibility and interpretation of hippocampal research in health and disease.

## 2. Materials & methods

### 2.1. Subject recruitment

A total of 44 healthy subjects were included in this study. Subjects were recruited from two separate acquisition sites after providing written informed consent in accordance with the Declaration of Helsinki. For both acquisition sites, ethical approval for the experimental procedures was provided by their institutional ethics review boards (i.e., Faculty of Psychology and Neuroscience, Maastricht University, the Netherlands, and Health Sciences Research Ethics Board of Western University, London, Canada), respectively. For the following sections, we refer to these datasets as the ‘Maastricht’ (N=16, age = 38.40 ± 14.24, between 20 and 66 years old, 4 males) and ‘London’ (N=28, age = 46.14 ± 12.84, between 20 and 66 years old, 18 males) dataset.

### 2.2. MRI acquisition

MR images from both acquisition sites were acquired on a Siemens 7T scanner (Siemens Healthineers, Erlangen, Germany), but differed in their gradient system type (i.e., head-only vs. whole-body), as well as RF head coil (see Table 1). Sub-millimeter MP2RAGE anatomical (0.7 mm isotropic nominal voxel size), as well as lower resolution Sa2RAGE (2 mm isotropic nominal voxel size) data were acquired to quantify T_1_ and map B_1_^+^ (see Figure 1A for an example B_1_^+^ map for each acquisition site) across the brain. The time resampled frequency offset compensated inversion (TR-FOCI) pulse was implemented for the MP2RAGE sequence at both acquisition sites to improve inversion efficiency and T_1_ quantification (Hurley et al., 2010). See Table 1 for further details on the acquisition set up and sequence parameters.

**Table 1.**
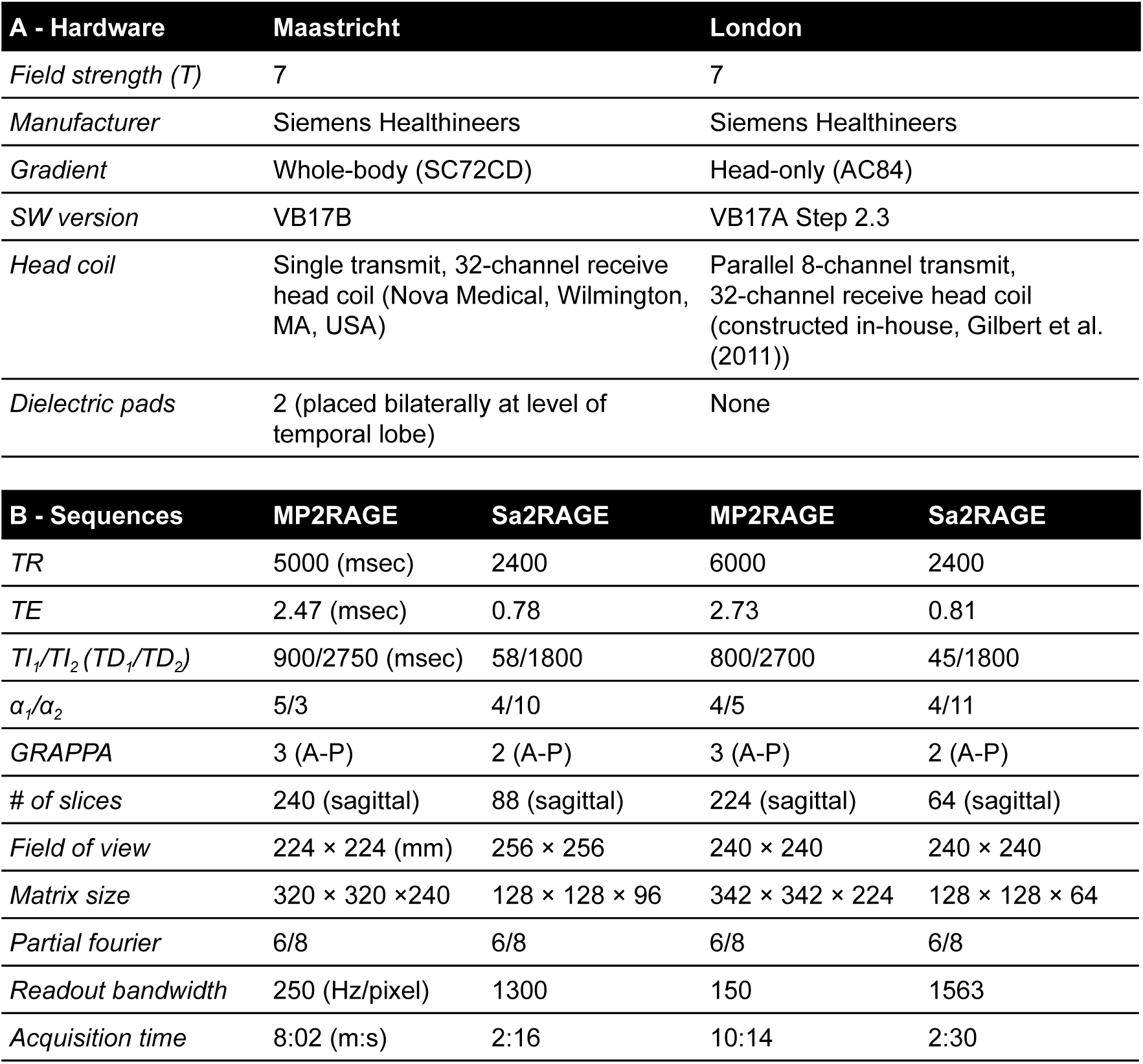
MRI scanner and sequence set ups, specified per dataset.

**Figure 1.**
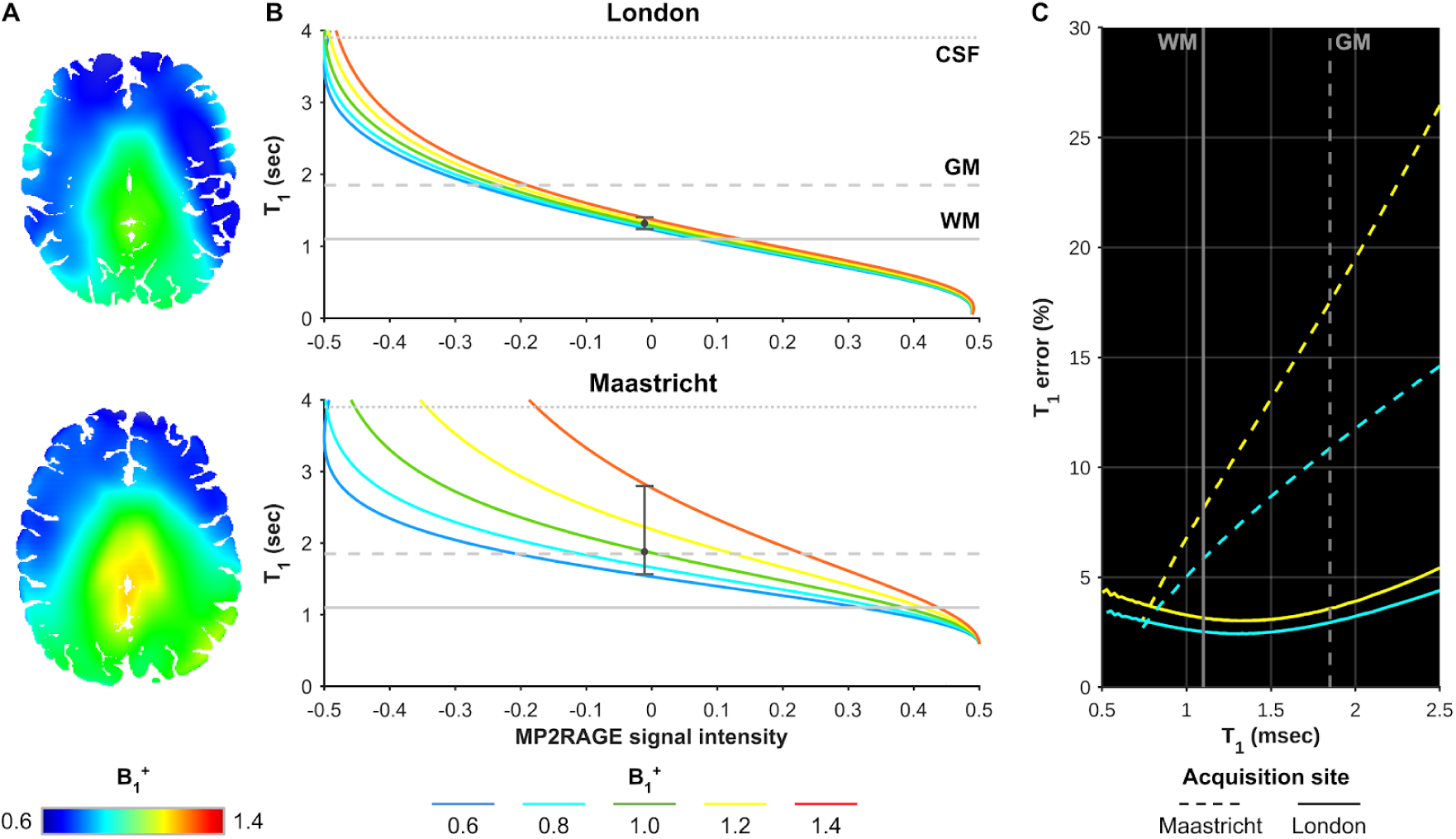
MP2RAGE B ^+^ dependency. (A) Example B_1_^+^ map for each acquisition site. (B) B_1_^+^ dependency of the T_1_ map for a range of B_1_^+^ values (colored solid lines) for each acquisition site’s MP2RAGE protocol (top and bottom panel). Typical WM, GM, and CSF T_1_ values are indicated using the vertical lines. (B) Differences between acquisition sites (dashed vs. solid lines) in the calculated T_1_, as a function of T_1_, due to B_1_^+^ values that are ±20% different (i.e., 0.8 vs. 1.2, in cyan and yellow, respectively) from the nominal value. See section 4.3. for a discussion on potential factors underlying this inter-site difference in B_1_^+^ sensitivity.

For the pTx system, mapping of the default excitation mode (B_1_^+^) was performed with Actual Flip Angle Imaging (AFI) with optimized RF and gradient spoiling (flip-angle: 70°, TR_1_/TR_2_: 30/150 ms, resolution: 3.75 mm isotropic, matrix: 64 × 64 × 48, orientation: sagittal, partial Fourier sampling: 6/8 in both phase encoding directions, Yarnykh (2007) and Nehrke (2009)). This was complemented by low flip-angle GRE images of the same geometry using the Fourier encoding scheme with a TR of 7 ms in order to generate absolute calibrated flip angle maps (Brunner and Pruessmann, 2009; Nehrke and Börnert, 2008; Tse et al., 2014). To simultaneously obtain a B_0_ map, the AFI sequence was acquired with five echoes (echo-times: 1.9, 3.4, 4.9, 6.3, and 7.8 ms, total acquisition time: 5 min). Shimming of the transmit field was accomplished using a magnitude least squares optimization of the field intensity over the specified adjustment volume using the calibrated flip angle maps and optimizing using only the phase of each transmit channel. For the single channel system, B_0_ maps were acquired during the regular Siemens prescan with two echoes.

### 2.3. Data pre-processing

Before the B_1_^+^ correction, datasets from both acquisition sites were pre-processed as described in Haast et al. (2018b). This included brain extraction using an optimized skull-stripping workflow, and coregistration of the Sa2RAGE to the MP2RAGE data (also part of Haast (2019)). After coregistration, MP2RAGE data were corrected for B_1_^+^ inhomogeneities as described in the original paper (Marques and Gruetter, 2013) resulting in ‘corrected’ T_1_w (i.e., UNI) and quantitative T_1_ maps. In the following sections, we will refer to this dataset as ‘corrected’, while ‘original’ data denotes the uncorrected dataset. The B_1_^+^ dependency plots of the T_1_ maps, as well as calculated T_1_ error (%) as a function of T_1_, for the current sequence parameters used by each acquisition site are displayed in Figure 1.

### 2.4. Cortical segmentation analysis

The pre-processed MP2RAGE T_1_w images were used as input for the sub-millimeter longitudinal processing workflow implemented in the FreeSurfer (v6.0, http://surfer.nmr.mgh.harvard.edu/) image analysis suite to obtain brain tissue segmentations and white matter (WM) and pial surfaces reconstructions (Dale et al., 1999; Reuter et al., 2012). Longitudinal analyses of the data were necessary to allow direct (i.e., vertex-by-vertex) comparison between the surface reconstructions and cortical thickness surface metric based on either the original or B_1_^+^-corrected MP2RAGE T_1_w images. See Reuter and Fishl (2011) for more details. Reconstructed cortical surfaces for both acquisition sites were processed as described in the ‘postprocessing pipeline’ in Haast et al. (2018b) to quantify cortical T_1_ and thickness differences by comparing: (1) original vs. corrected dataset, as well as (2) Maastricht vs. London acquisition sites.

### 2.5. Subcortical segmentation analysis

In addition, FreeSurfer’s subcortical (i.e., ‘aseg’) output (Fischl et al., 2002), pooled from both acquisition sites, were used to study segmentation differences between original and corrected data, as well as acquisition sites and hemispheres, for the hippocampus as well as a given set of subcortical regions of interest (ROIs): thalamus, caudate nucleus, putamen, globus pallidus and nucleus accumbens. In addition to the aseg output, we also included the labels obtained by running the automatic hippocampal subfield segmentation implemented in FreeSurfer (Iglesias et al., 2015). Segmentations for each of the ROIs were compared before and after B_1_^+^ correction. This was based on the volumetric labels using total volume (in mm^3^), label overlap (i.e., Dice) and distance between label boundaries (i.e., Hausdorff distance) as in Gulban et al. (2018), and in surface space, based on shape differences using large deformation diffeomorphic metric mapping (LDDMM) as in Khan et al. (2019). See also Figure 2 for a schematic overview of the processing workflow.

**Figure 2.**
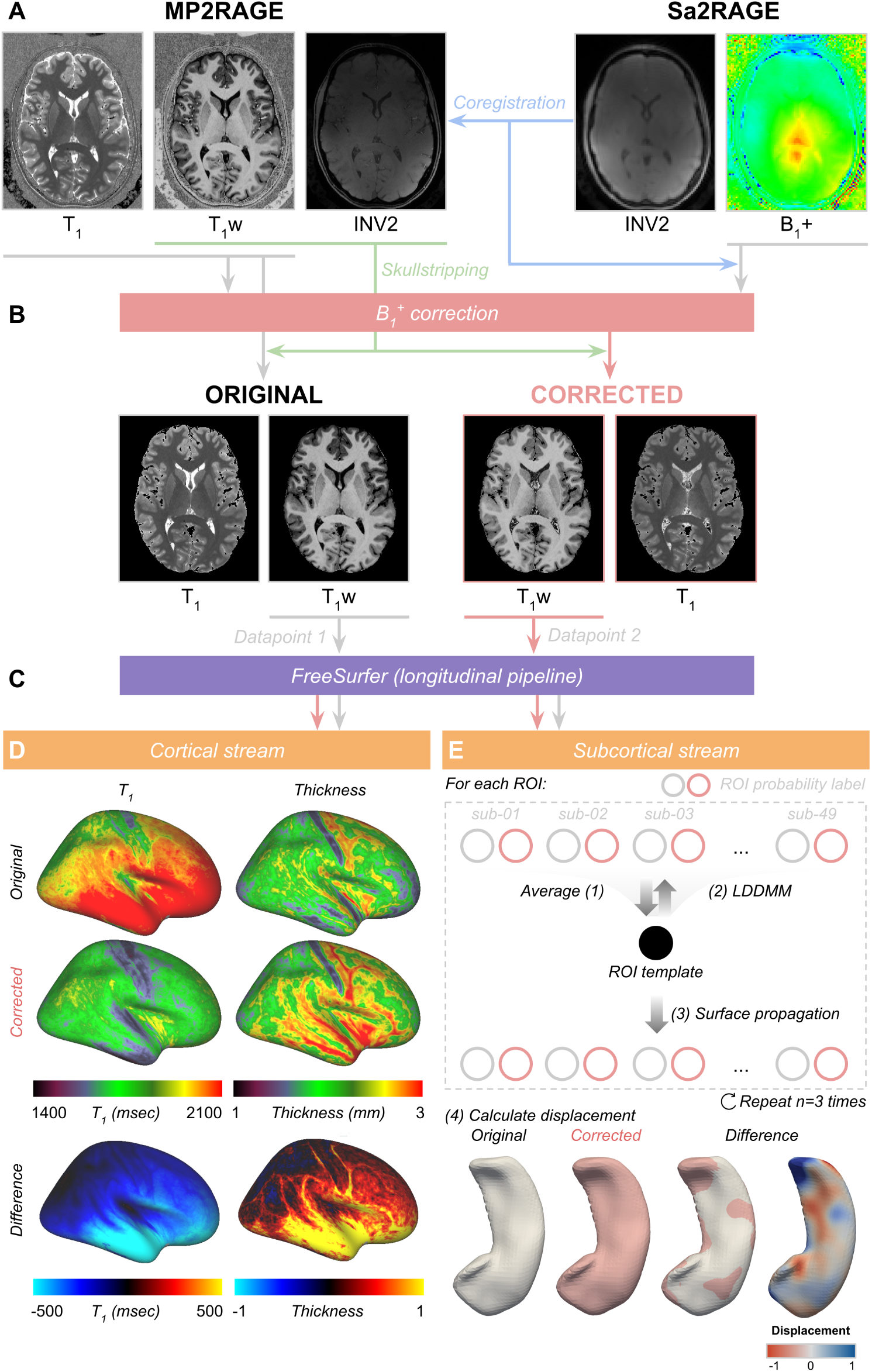
Cortical and subcortical analysis pipeline. For each subject, MP2RAGE T_1_w and T_1_ maps (A) were corrected for B_1_^+^ homogeneities using the coregistered Sa2RAGE B_1_^+^ map following the procedure described in Marques and Gruetter (2013) (B). Skull-stripped original and corrected T_1_w volumes were then used as a single data (i.e., ‘time’) point for FreeSurfer’s longitudinal analysis pipeline to reconstruct cortical surfaces with matching topology (C). Differences in cortical T_1_ and thickness between original and corrected datasets were calculated as described in Haast et al. (2018b) (D). In addition, morphometry was performed using the LDDMM algorithm (Beg et al., 2005; Khan et al., 2019) to quantify differences in subcortical segmentation after B_1_^+^ correction for each ROI (E).

#### 2.5.1. Volumetric assessment

First, we estimated for each subject within the Maastricht and London datasets their expected subcortical volumes, based on age, gender, estimated total intracranial volume and the scanner characteristics using the model presented in Potvin et al. (2016). Their averages were then used as estimates for FreeSurfer’s subcortical output. Second, to assess the correspondence between segmentation labels in terms of global shape and boundaries, we used the Dice coefficient and Hausdorff distance, respectively. The Dice coefficient is a common metric to quantify volumetric correspondence between two ROI segmentations – the original and corrected data, in our case – and a Dice score of 1 indicates perfect overlap (Taha and Hanbury, 2015). The Hausdorff distance score is a distance metric sensitive to boundary errors and thus can be used to quantify the similarity between the two boundaries. Here, a Hausdorff distance represents the average number of voxels by which the two boundaries deviate from one another (Taha and Hanbury, 2015). Both metrics have been used as implemented in the Nilearn package (v.0.5.0, Abraham et al. (2014))

#### 2.5.2. Surface-based assessment

Surface-based comparisons were performed following the procedure described previously (Khan et al., 2019) and using openly available image processing scripts developed in-house (https://github.com/khanlab/surfmorph). Fuzzy labels for each of the subject’s ROIs were obtained by smoothing the binary ROI label with a 1 × 1 × 1 mm kernel size. Here, left and right hemispheres were combined into a single volume and treated as a single label. The fuzzy (i.e., smoothed) labels for each of the ROIs, for each subject, were transformed to MNI space (Fonov et al., 2011) using linear transformations based on the subject’s corrected MP2RAGE T_1_w volume to MNI volume transformation. As for FreeSurfer’s longitudinal workflow, these linearly aligned labels were used to generate unbiased averages for surface generation. These averages were computed by iterating through steps of (1) template generation by averaging across subjects, and (2) registration of each segmentation image to this template using LDDMM registration (Beg et al., 2005). The resulting fuzzy segmentation was then used to generate the ROI’s template surface through a 50% probability isosurface. The 3D volume of the ROI’s template was then fit to each subject’s segmentation using LDDMM, with affine initialization to provide vertex-wise correspondence between all surfaces of that specific ROI. The template surface was then propagated to each subject’s ROI, to provide surfaces with common indices for performing vertex-by-vertex morphometry analyses and mapping of tissue contrast near the the vertex’ positions.

#### 2.5.3. Generation of subcortical surface maps

First, to allow shape analyses, in-/outward displacements at each vertex location were computed between the template surface and the injected subject surface, using the projection along the surface normal. Importantly, the mean displacement across a spherical neighbourhood (10 mm radius) was computed for each vertex and subtracted from the local vertex-wise displacement. This effectively ensures local displacements are not affected by residual positional differences that could remain after the linear alignment between template and subject.

Second, to obtain rough estimates of contrast changes near the ROI boundaries that may affect automatic segmentation, and thus surface placement, gradient magnitude maps were computed for the original and corrected FreeSurfer white matter normalized (i.e., ‘T1.mgz’) input. This was done using the ‘*-volume-gradient*’ function within the Connectome Workbench command-line tool (Marcus et al., 2011). The local change in gradient magnitude (i.e., corrected−original gradient maps) were then sampled at the original vertices’ coordinates and smoothed using the structure’s surface geometry (i.e., across neighbors). Resulting maps were added as scalar data to the surfaces meshes VTK files for visualization and vertex-wise analyses.

Finally, the minimal geometrical distance (in mm) for each vertex on the ROI’s template surface to the CSF based on MNI’s CSF probability map was calculated for follow-up analyses.

### 2.6. Statistical analysis

Repeated measures analysis of variance (ANOVA) was used to compare subject-averaged T_1_ and thickness values between acquisition sites (between-subjects factor), as well as to test for a B_1_^+^ correction effect (i.e., original vs. corrected: within-subject). For hippocampal analyses, we used a mixed model ANOVA to test the main effect of B_1_^+^ correction (within-subject) on volume (in mm^3^) and related volumetric metrics (i.e., Dice coefficient and Hausdorff distance). Potential differences between hemispheres (within-subject), acquisition sites (between-subjects) and/or interactions with the main effect were statistically tested by including them in the mixed ANOVA model.

Differences in hippocampal and subcortical surface displacement and changes in gradient magnitude, as result of the B_1_^+^ correction, were compared between acquisition sites using the SurfStat toolbox (http://www.math.mcgill.ca/keith/surfstat/) for Matlab (R2018b, The Mathworks, Natick, MA, USA). Here, vertex-wise one-sided T-tests were used for identification of vertices characterized by larger absolute surface displacement, and gradient change for the Maastricht dataset. Resulting statistical maps were corrected for multiple comparisons using random field theory for non-isotropic images (Worsley et al., 1999) and mapped as scalar data to the surface meshes VTK files.

## 3. Results

### 3.1. Inter-site comparison of cortical T_1_ and thickness

Site-averaged cortical T_1_ and thickness data are displayed in Figure 3A and B, respectively. Identical data scaling across datasets and within data modalities were used as much as possible for inter-site comparison purposes. A pronounced discrepancy in T_1_ can be observed between the Maastricht and London data sets (i.e., average cortical T_1_ of 1967.56 vs 1701.49 ms, respectively, *F*(1,41)=334.68, *p*<.001) based on both the cortical pattern (Figure 3A), as well as the corresponding histograms (Figure 4A). A similar trend is visible for cortical thickness with lower average cortical thickness seen in the Maastricht data (2.11 vs 2.26 mm, *F*(1,41)=57.58, *p*<.001).

**Figure 3.**
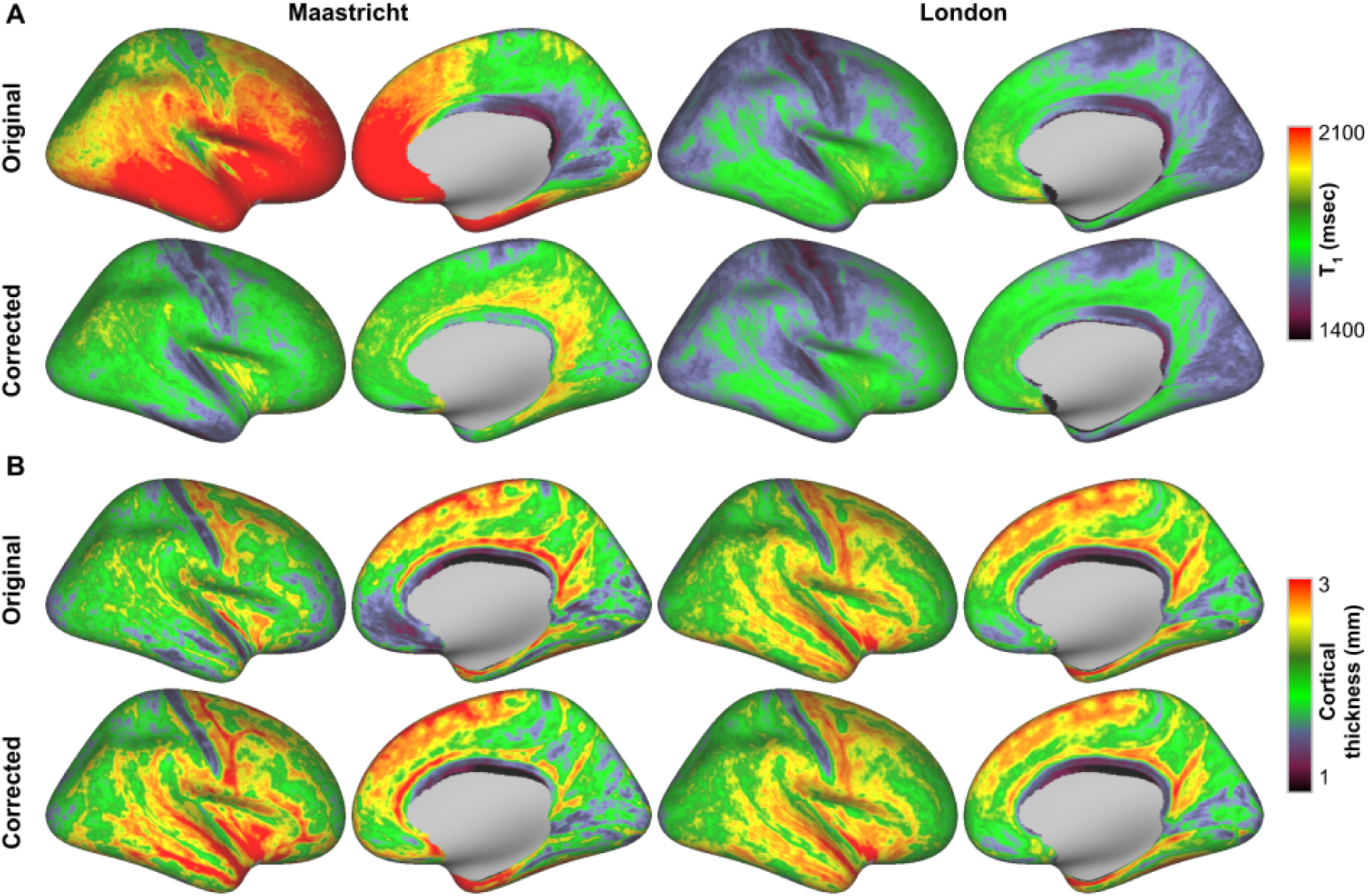
Average cortical T_1_ and thickness surface maps. Original (odd rows) and corrected (even rows) T_1_ (ms, A) and thickness (B) were mapped onto an inflated right hemisphere surface and averaged across all subjects. Left (Maastricht) and right (London) columns represent different acquisition sites. Data is scaled identically within data modality for comparison.

**Figure 4.**
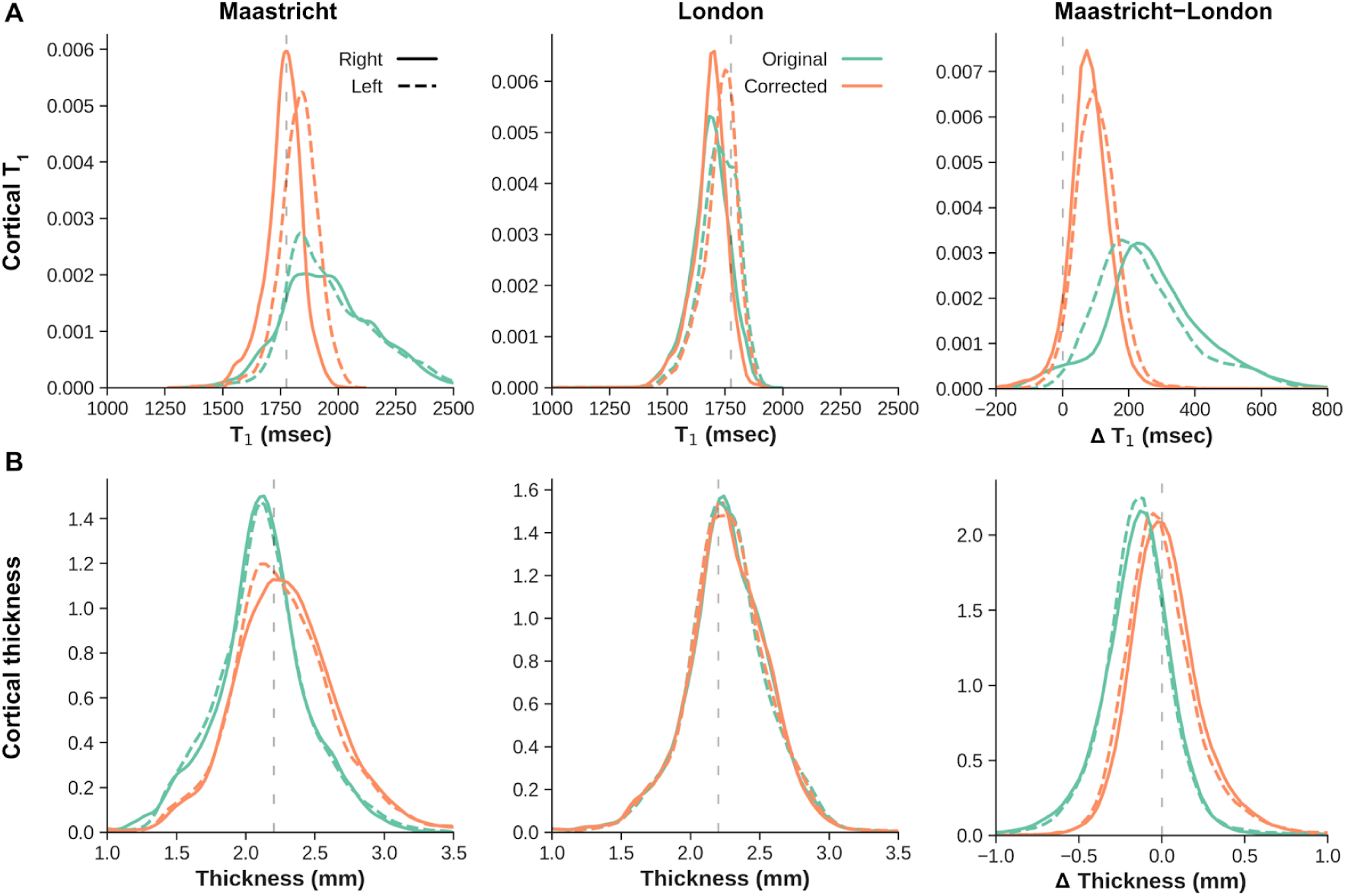
Cortical T_1_ and thickness distribution. (A) T_1_ (ms) and (B) thickness distributions before (green) and after (orange) B_1_^+^ correction. Histograms are shown for both right and left hemispheres (solid and dashed linestyle, respectively), acquisition sites (left and middle columns) and inter-site difference (right column). Vertical dashed gray lines are shown for comparison

Figure 4 demonstrates the increased similarity between acquisition sites, improving from an average inter-site T_1_ differences (i.e., Maastricht−London, top right panel) of 266.04 ms before to 94.11 ms after B_1_^+^ correction. For cortical thickness, the difference decreased from −0.14 mm to 0.02 mm (bottom right panel). This difference drop predominantly originates from a significantly stronger change for the Maastricht data based on a site*correction interaction effect on average cortical T_1_ (*F*(1,41)=75.58, *p*<.001) and thickness (*F*(1,41)=177.19, *p*<.001) values. Initially, clear biases with higher T_1_ and lower cortical thickness in the temporal and frontal lobes (i.e., low B_1_^+^ regions, see Supplementary Figures 1 and 2) are observed in the original Maastricht data (Figure 3). However, differences between acquisition sites become more homogeneous, and centered more closely around 0 msec / 0 mm, after correcting the MP2RAGE data for B_1_^+^ inhomogeneities based on the histograms.

### 3.2. The effect of B_1_^+^ correction on automatic hippocampal segmentation

Figure 5 shows a comparison of the hippocampal aseg labels after running the original and corrected MP2RAGE T_1_w image through the longitudinal FreeSurfer pipeline. Visual inspection of the labels’ reconstructed surface meshes (Figure 5A) shows local differences in surface placement between the original and corrected labels (i.e., yellow vs. red). Please, note that surfaces are taken from a single subject within the Maastricht dataset, as – in line with the observations in the cortex – these subjects were characterized by larger differences in subcortical volumes as well. In this example, clear examples of in- and outward movement of label boundaries are indicated using dotted and solid arrows, respectively. In general, we observe the strongest changes near the hippocampal tail and head regions. We quantified the label overlap and label boundary distances using the Dice and Hausdorff scores for all subjects (see Figure 5B). These reveal significantly larger overlap (0.97±0.01 vs 0.90±0.03, *F*(1,42)=266.41, *p*<.001), but smaller boundary distances (2.75±1.13 vs 3.88±0.78, *F*(1,42)=23.20, *p*<.001) between original and corrected labels for the London data. Also, we observe that changes after B_1_^+^ correction are stronger for the right hemisphere labels (*F*(1,42)=43.04, *p*<0.001), as indicated by the lower Dice coefficient scores compared to the left hemisphere across both acquisition sites (0.88±0.03 vs 0.91±0.02 and 0.96±0.01 vs 0.98±0.01 for Maastricht and London, respectively). Similar trends are seen based on the average Hausdorff distances between label boundaries (*F*(1,42)=7.78, *p*<.01).

**Figure 5.**
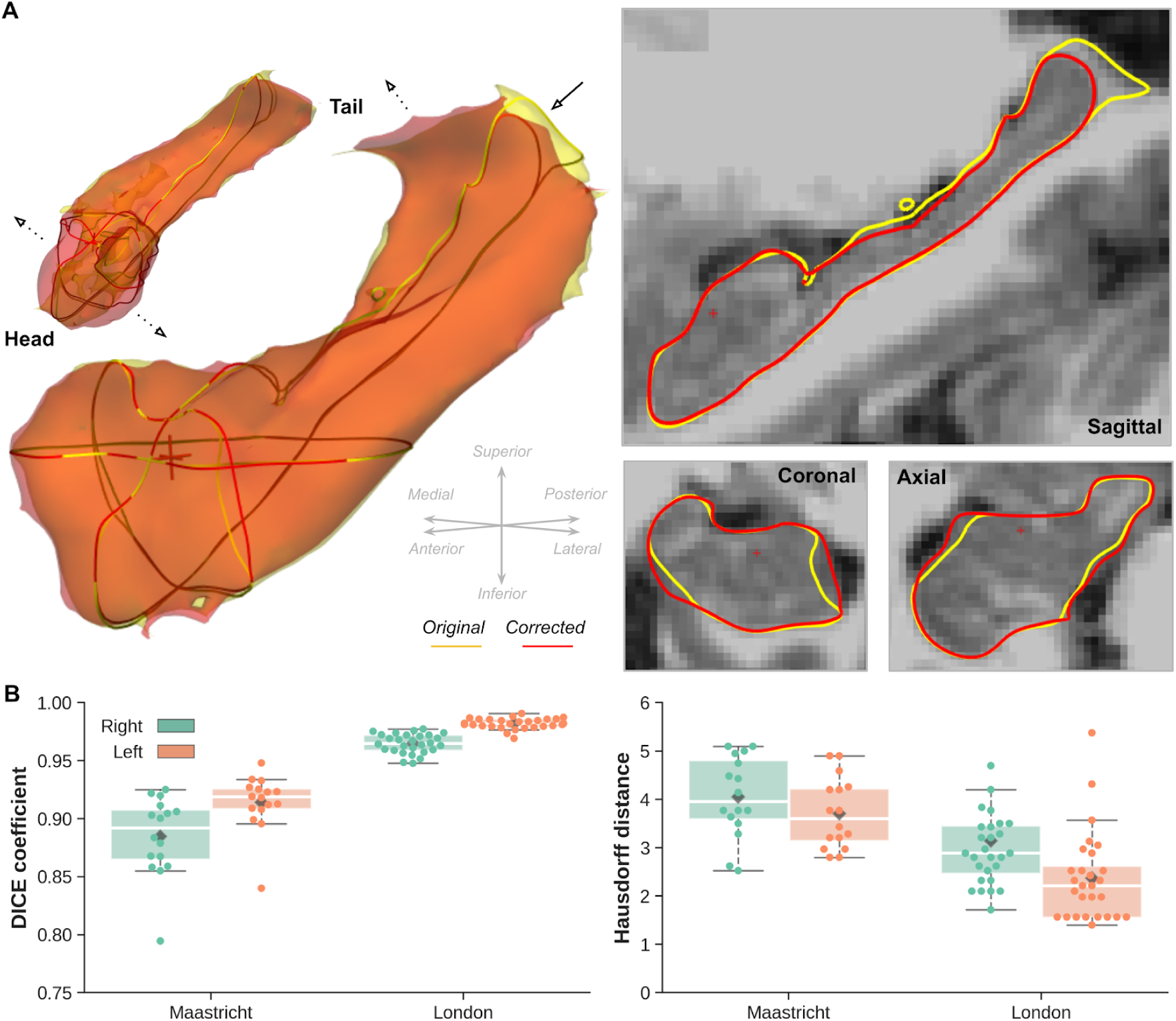
Hippocampal segmentation. (A) Single-subject example of reconstructed left and right hemisphere hippocampal surface meshes after processing the original (yellow) and corrected (red) MP2RAGE data. Solid and dotted arrows indicate clear examples of in-/outward movement of label boundaries after B_1_^+^ correction, respectively. Corresponding sagittal, coronal and axial cross sections of the left hemisphere surface outlines are shown on the right overlaid onto the corrected MP2RAGE T_1_w map. (B) Box plots showing distribution of original vs. corrected hippocampal Dice coefficient (left) and Hausdorff distance (right) for both acquisition sites (x-axis), and right (green) and left (orange) hemispheres. Box and whisker extends demarcate interquartile ranges and distribution (excluding outliers), respectively, while diamonds and dots represent group means and individual subjects data, respectively.

Figure 6 quantifies the ‘global’ differences in volume (in mm^3^, A), between hippocampal aseg labels before and after B_1_^+^ correction (B), and between hemispheres (C). Averaged across hemispheres, slightly lower hippocampal volumes (main acquisition site effect, *F*(1,42)=3.22, *p*=.08) are observed for the Maastricht dataset (3419.51±289.33 mm^3^) compared to the London dataset (3603.74±363.82 mm^3^), which align more closely with the ‘expected’^1^ volumes (horizontal dashed lines at 3641.46±156.52 mm^3^, Figure 6A). In line with the changes in cortical thickness, B_1_^+^ correction has a smaller effect on global hippocampal volume in the London dataset (+52.54±50.26 mm^3^). Slightly larger changes are observed for the Maastricht data (+80.15±208.13 mm^3^), especially after taking into account differences across left (−37.94±108.14 vs 24.33±23.59 mm^3^ for Maastricht and London, respectively) and right hemispheres (+198.25±216.91 vs 80.75±53.89 mm^3^) as indicated by a significant interaction effect (F(1,42)=18.83, p<.001), see Figure 6B. This relates to the differences in the Dice and Hausdorff scores (see Figure 5B). As a result, inter-hemispheric differences become more comparable across acquisition sites (Figure 6C). See Supplementary Data 1 for evaluation of the B_1_^+^ effects on automatic hippocampal subfields segmentation.

**Figure 6.**
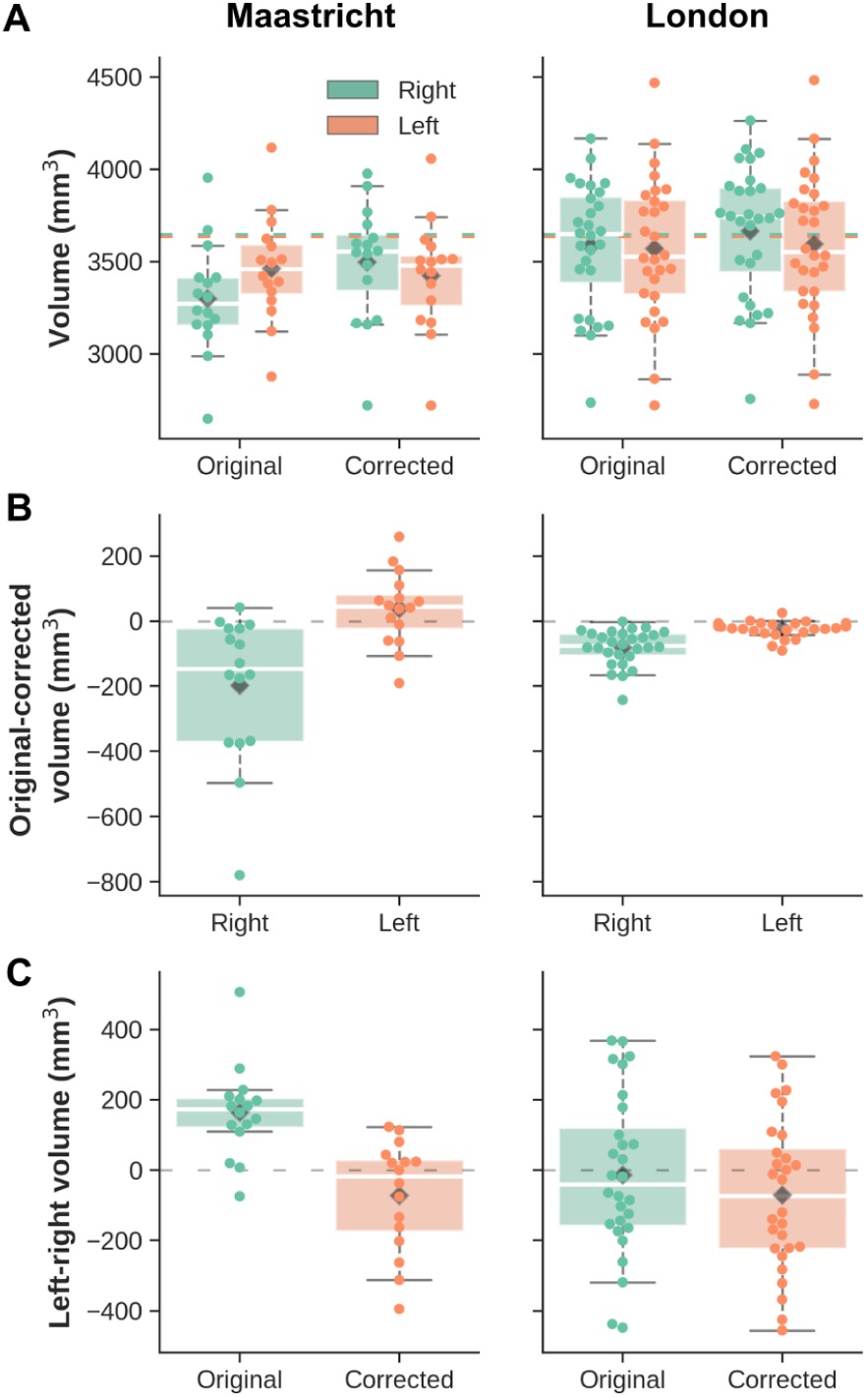
Hippocampal volume. Box plots for each acquisition site (left and right column) showing (A) left (orange) and right (green) hemisphere hippocampal volume (in mm^3^) before and after B_1_^+^-correction and corresponding estimates (horizontal dashed lines), (B) original−corrected difference in hippocampal volume for right (green) and left (orange) hemispheres, and (C) left−right difference in hippocampal volume before (green) and after (orange) B_1_^+^ correction. Box and whisker extends demarcate interquartile ranges and distribution (excluding outliers), respectively, while diamonds and dots represent group means and individual subjects data, respectively.

While the data shown so far allowed us to compare the global changes in hippocampal volume after B_1_^+^ correction, between hemispheres and acquisition sites, the surface displacement measures shown in Figure 7 allow us to quantify and localize the changes in label boundaries placement more precisely. Figure 7A shows the same single-subject example from the Maastricht dataset. Here, in line with the observations in the volume space (see Figure 5A), surface placement changes more strongly closer to the tail and head regions, as apparent by the deep blue and red coloring. These indicate in- & outward placement of the boundaries in the original data, respectively. In addition, as these surface displacements were computed in MNI space, we were able to perform group comparisons. After averaging across subjects for each acquisition site, the extent of surface displacement is clearly larger in the Maastricht dataset (s.d.=0.10, across both hemispheres, left column) compared to the London dataset (0.01, right), with averages both centering around 0 (Figure 7B). Distributions for left and right hemispheres are shown using solid and dashed lines, respectively. Especially for the latter, the hippocampal surfaces were positioned more inwards for the original data. Inset figures show the corresponding sagittal cross sections of the average differences in displacement along the surface mesh overlaid onto the MNI template. Please note that a different scaling (×10 difference) was used between acquisition sites in order to appreciate their patterns. In contrast, similar scaling was used for the surface maps shown in Figure 7C, from both an anterior (top row) and posterior (bottom) perspective. Dotted patterns indicate the vertices which were characterized by significant larger surface displacement (*p*<.05, multiple-comparison-corrected) for the Maastricht dataset compared to the London dataset. Again, these tend to localize more towards the tail and head regions, in line with the observations for the single subject data shown in Figure 5A.

**Figure 7.**
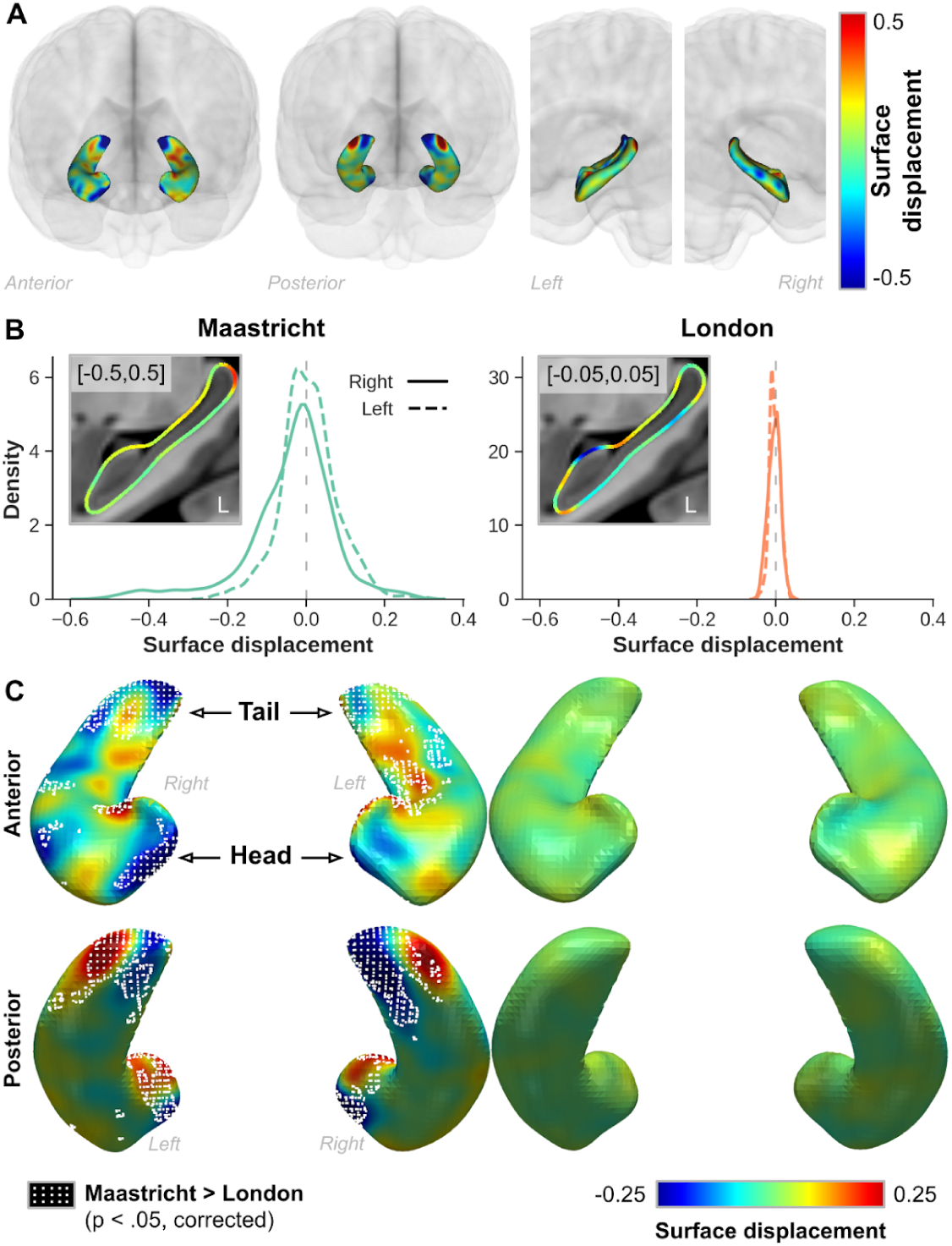
Hippocampal displacement. (A) Single-subject example from the Maastricht dataset showing the color-coded hippocampal surface displacement between original and corrected data. (B) Distribution plots showing the averaged hippocampal surface displacement for the Maastricht (left) and London (right) datasets. Solid and dashed lines indicate right and left hippocampal data, respectively. Inset figures show corresponding sagittal cross section of the left hemisphere hippocampus. (C) Surface representation of average displacement for both acquisition sites shown from an anterior (top) and posterior (bottom) perspective. Regions where the Maastricht data was characterized by significantly (p < .05, multiple comparison corrected) larger surface displacement are demarcated using a white dotted pattern. In all cases, blue and red colored surface displacement indicate regions where label boundaries were placed more in- and outwards, respectively, in the original data.

As we found changes in surface placement due to the B_1_^+^ correction (Figure 7), we mapped the change in gradient magnitude at the vertices coordinates onto the surface meshes to detect whether changes in tissue contrast colocalize with changes in surface displacement. First, Supplementary Figure 3A highlights the distributions of changes in the gradient for both acquisition sites. Again, the Maastricht data is characterized by a wider distribution (*s.d.*=2.94 vs. 0.62) with larger changes occurring at the right hippocampus (solid lines). Statistical testing reveals that the differences between acquisition sites are spatially widespread, as indicated by the white dotted pattern, but tend to localize more towards the lateral (i.e. ‘outside’) and longitudinal extents (i.e., head and tail, Supplementary Figure 3B). The tail region seems to be affected most considering the overlap (see purple patches in Supplementary Figure 3C) of the statistical maps based on both surface displacement (red), as well as gradient change (blue).

Next, the volumetric and 3D representations in Supplementary Figure 4A and B, respectively, allow us to better understand the spatial pattern observed for the changes in gradient magnitude. For the same subject, used as example in Figures 5A and 7A, we observe the strongest changes in surface placement (i.e., based on the misalignment of the original, yellow, and corrected, red, label boundaries in Supplementary Figure 4A) and/or gradient magnitude (i.e., color scaled surface mesh in Supplementary Figure 4C) closer to bordering structures including CSF and arteries, demarcated by the cyan and red arrows/meshes, respectively. As such, we reasoned that surface displacement and/or change in gradient may correlate with the distance to CSF (see Supplementary Figure 5), which is characterized by strong changes in intensity due to the B_1_^+^-bias removal procedure (see Figure 1A). The top plot in Supplementary Figure 4C (hippocampal data only) does not show a direct relationship between surface displacement averaged across subjects and distance (orange line) to CSF (x-axis), but does reveal that the gradient changes (scatter plot) become less strong moving away from CSF. In addition, we observe that changes in the vertices’ placements are more variable across subjects (green line) the closer it is to CSF. This pattern is consistent (bottom plot in Supplementary Figure 4C) when including data from the remaining subcortical structures.

For the analyses of volume and surface placement changes of the subcortical structures, we refer to Supplementary Figure 6. Briefly, in line with the hippocampal results, we observe a lower average subcortical volumes (A), as well as stronger changes in volume after B_1_^+^-correction (B) for the Maastricht dataset. In general, inter-hemispheric differences become more comparable across acquisition sites (C). As a result, surface displacement varies significantly more (dotted pattern in C, p<.05, corrected) for the Maastricht data (left column) compared to the London dataset (right), with more variable changes in surface placement closer to the CSF (see Figure 9C, bottom plot).

<u>Link to Supplementary Figure 6</u>

## 4. Discussion

We have shown before that B_1_^+^ residuals in MP2RAGE data affect performance of the brain’s cortical analyses by means of cortical T_1_ and thickness biases (Haast et al., 2018b). Cortical T_1_ values were artificially high, and thickness estimates too low in regions characterized by low B_1_ ^+^, and vice versa in regions with high B_1_^+^. However, as advocated in the original MP2RAGE papers (Marques et al. (2010); Marques and Gruetter (2013)), B_1_^+^ sensitivity – that is, the degree of B_1_^+^-related image inhomogeneity that still resides in your MP2RAGE data – greatly depends on your sequence setup, and scanner hardware. As such, the work presented in the current paper validated this dependency by extending our analyses to an independent dataset acquired at a different 7T MRI site (CFMM, London, ON, Canada). Here, MP2RAGE data were (1) acquired using sequence parameters that rendered the images minimally B_1_^+^ sensitive; and (2) combined with favorable MR hardware for achieving increased B_1_^+^ homogeneity.

### 4.1. Inter-site cortical T_1_ and thickness variability

Our results show substantial non-biological variability in cortical T_1_, as a result of differences in B_1_^+^ sensitivity and B_1_^+^ field homogeneity, between acquisition sites. These discrepancies in T_1_ (and underlying MP2RAGE signal intensities) translate towards strong local differences in cortical thickness measures, especially in the critical brain regions such as temporal and frontal lobes which are typically characterized by strong B_1_^+^ offsets at 7T (see Supplementary Figures). Most importantly, while striking differences between acquisition sites (Maastricht−London) were visible before B_1_^+^ correction in terms of cortical T_1_ (i.e., 266.05 ms difference on average) and thickness (−0.14 mm), these were substantially equalized (94.11 ms and 0.02 mm, respectively) after removing the B_1_^+^ bias. These insights are of high importance when comparing or pooling MP2RAGE-based cortical T_1_ and thickness data between or across subjects acquired as part of different studies and/or at different sites. Note that this is not only true for the MP2RAGE sequence but would apply for other sequences, based on the same principle – i.e., acquisition of different steady-state conditions using varying excitation angles – as well (Deoni et al., 2004; Venkatesan et al., 1998). In terms of cortical longitudinal relaxation times, we observed an average cortical T_1_ (or longitudinal relaxation rate 1/T_1_) of ∼1750 ms (0.57 s^-1^) across both data sets. This is slightly lower than the ∼1900 ms (0.53 s^-1^) in the motor and temporal cortices measured at 7T by Marques et al. (2010), but closer to those observed by Metere et al. (2017). Interestingly, we detected a similar offset between left and right hemispheric T_1_ across acquisition sites, with higher values observed for the left hemisphere. This follows the inter-hemispheric differences with lower R_1_ (i.e., higher T_1_) values for the left hemisphere observed by Shams et al. (2019), across different field strengths (Kim et al., 1994; Marques et al., 2017; Wansapura et al., 1999), and which could not solely be explained by the asymmetric B_1_^+^ field. In fact, our results show that this difference intensifies after cleaning up remaining B_1_^+^ residuals. Learning- and aging-induced cortical myelination changes are not unknown and could lead to region-specific T_1_ differences between hemispheres (Callaghan et al., 2014; Natu et al., 2019). It is worth noting that these differences in T_1_ do not extrapolate towards hemispheric cortical thickness differences. Further research, e.g. by disentangling the relationship of the subject’s functional lateralization of the brain and handedness on regional T_1_ (Toga and Thompson, 2003), would be necessary to investigate this apparently more systematic inter-hemispheric T_1_ bias. However, the effect of handedness may be negligible considering the fact that all subjects within the London dataset were right-handed while an inter-hemispheric difference in T_1_ was still visible.

### 4.2. Effect of B_1_^+^ inhomogeneities on hippocampal morphometry

Besides the effects on cortical T_1_ and thickness measurements, residual B_1_^+^ inhomogeneities may also affect performance of hippocampal and subcortical segmentations. Accurate segmentation of brain structures such as the hippocampus, caudate nucleus and putamen are of essence for neurodegenerative research, which heavily relies on accurate assessment of these structure’s sizes across their patient populations. In practice, segmentation of the subcortical structures is often achieved by deforming the subject’s anatomical scan to a common atlas (Keuken and Forstmann, 2015; Mazziotta et al., 2001; Xiao et al., 2017) or using voxel-based neuroanatomical labeling tools (Bazin et al., 2014; Fischl et al., 2002). However, the former becomes problematic in case the study population deviates from the population used for generating the atlas (i.e., patients vs. healthy controls). In these cases, use of segmentation algorithms to label the different tissue types based on the subject’s anatomical (e.g. T_1_w) image(s) is preferred. However, as we observed for the cortical gray matter, residual B_1_^+^ inhomogeneities can substantially affect image contrast and may therefore introduce variability in the performance of automated methods to precisely define the regions and their borders, which may affect study replicability. Here, we used volume- and shape-based analyses to examine the extent of a potential B_1_^+^-related bias on non-neocortical segmentation within and across acquisition sites, with a major focus on the hippocampus. Structural changes of the hippocampus, such as reduced volume (i.e. atrophy), are well-established biomarkers in numerous neurodegenerative and psychiatric diseases linked to memory loss (Small et al., 2011). Previously it was shown that proton density (i.e. B_1_^-^) effects in MPRAGE T_1_w data modulates subcortical results, especially in regions where the GM-WM contrast is low. Small artificial deviations in the accuracy of subcortical boundary definitions led to spurious brain morphological changes (Lorio et al., 2016). As such, similar methodological-related biases, in the form of image inhomogeneities related to B_1_^+^, or due to use of different segmentation tools vs. manual tracing (Morey et al., 2009; Wenger et al., 2014) may introduce artificial variability across individuals surpassing the volume differences observed between normal and diseased populations (Lupien et al., 2007).

While the output of FreeSurfer’s subcortical segmentation from the London dataset were relatively stable, hippocampal volumes changed significantly after removal of residual B_1_^+^ inhomogeneities from Maastricht’s MP2RAGE data. As a result, estimated hippocampal volumes shifted more closely towards the expected range of hippocampal volumes, based on (1) the averages obtained using the model by Potvin et al. (2016); (2) earlier observations across a wide age-range and gender-mixed population of healthy subjects (Lupien et al., 2007); and (3) the decreased inter-site difference, falling in line with the cortical thickness observations. Moreover, we observed a bias towards stronger hippocampal volume changes, predominantly increases, in the right hemisphere leading to a comparable left-right volume ratio across acquisition sites, i.e., characterized by slightly larger right hippocampi. This finding could possibly be attributed to the fact that the right hemisphere is usually larger than the left in right-handed individuals, and leads to a larger hippocampus as such (Xu et al., 2000). The hippocampus, however, is not a single uniform structure, but is composed of several components, or subfields, that are significantly distinct in terms of their cytoarchitectonic, vascular, and electrophysiological properties (Duvernoy, 1988). More advanced segmentation and/or optimization procedures, respecting these different hippocampal subfields, would therefore be beneficial to improve segmentation robustness. Indeed, optimization of the hippocampal labels using Bayesian inference and a statistical atlas within FreeSurfer’s automated hippocampal subfield segmentation tool (Iglesias et al., 2015) reduced the effect of B_1_^+^-related inhomogeneities on segmentation performance (see Supplementary Data) based on the improved Hausdorff distance scores between original and corrected MP2RAGE datasets.

We performed surface-based analyses by computing shape- and tissue contrast (i.e., gradient)-based metrics to localize and characterize these volume changes more precisely. In line with the inter-site differences in volume change due to the B_1_^+^ correction, the Maastricht dataset was characterized by more pronounced changes in hippocampal shape as well. Here, significant differences in shape between London and Maastricht were found to be located closer to the head and tail. Inter-site differences in hippocampal gray matter and white matter contrast were more widespread but tend to localize near structures characterized by most strongest T_1_ adjustments. Since changes were strongest for the Maastricht setup, we zoomed in on this dataset to more closely assess the relationship between the observed shape and contrast changes. As such, definition of the hippocampal boundary seems to become more variable (i.e., erroneous) in the original data near large arterial structures, or where the hippocampus is neighbored by thin strands of white matter and CSF interfaces. However, as illustrated in Figure 5A, white matter tissue near the hippocampal tail, or arterial voxels near the body are correctly excluded from the hippocampal label, while hippocampal gray matter is correctly included in the head after MP2RAGE correction. Although these findings do not warrant overall improvement in hippocampal segmentation accuracy by solely removing residual B_1_^+^ inhomogeneities, it does highlight the importance of careful consideration of the sequence parameters, taking into account the observed variability in hippocampal volume and shape due to B_1_^+^. In the following section, we will therefore compare both MP2RAGE setups used in this study and reiterate the importance of some of its parameters (Marques et al., 2010; Marques and Gruetter, 2013).

### 4.3. Effect of MP2RAGE parameters and MRI hardware

In case of the MP2RAGE sequence, the resulting T_1_ map is calculated using a lookup table based on the combination (UNI) image of two gradient-recalled echo datasets (GRE_1_ and GRE_2_) with different excitation angles (ɑ_1_ and ɑ_1_). However, due to the variable B_1_^+^ field, the flip angles can spatially vary, requiring post-hoc corrections to take this into account. Here, different combinations of parameters will affect the sensitivity of the UNI image to B_1_^+^ variations. This becomes already clear from the differences observed between the corresponding lookup tables, linked to the MP2RAGE acquisition setups (i.e., parameters and field strength), used for the voxel-wise estimation of T_1_ at each acquisition site. The increased sensitivity to B_1_^+^ differences across the brain for the Maastricht dataset is apparent based on the larger range of plausible T_1_ values per MP2RAGE signal intensity; and as a result, the actually observed cortical T_1_ values. This difference emphasizes the importance of B_1_^+^ correction for data acquired using this specific set of parameters as errors larger than 25% were not unusual. In line with earlier simulations by Marques et al. (2010), differences in B_1_^+^ sensitivity originate predominantly from the flip angle intensity and/or combination of flip angles for acquisition of the GRE datasets, and repetition time. While the London parameters follow the parameters recommended originally to be particularly B_1_^+^ insensitive, flip angles utilized in Maastricht were chosen mostly to emphasize intracortical tissue contrast due to variability in myelination (Geyer et al., 2011; Haast et al., 2016), and/or changes due to disease (Haast et al., 2018a). The flip angle during the first GRE image (ɑ_1_) has a large impact on the range of T_1_ values, especially towards the CSF spectrum of MP2RAGE signal intensities (i.e., left extent of Figure 1A). As the first flip angle is reduced (i.e., 4° instead of 5° in our case), the spread of T_1_ becomes more constricted (Marques et al., 2010). This additionally implies that the B_1_^+^ correction step would most strongly affect the CSF-GM tissues boundaries, which agrees with the (1) problematic delineation of the cortical GM and CSF tissue interface (Haast et al., 2018b); and (2) more variable subcortical boundary definitions closer towards CSF observed here. Moreover, the shorter repetition time (5000 vs. 6000 ms) but otherwise same nominal spatial resolution (0.7 mm^3^) will increase the B_1_^+^ sensitivity of the Maastricht setup due the increased number of excitations per MP2RAGE repetition time (Marques and Gruetter, 2013). Nevertheless, the exact effects of acquisition parameters on the sequence’s B_1_^+^ sensitivity may be more complicated than described above and should be considered carefully (Metere et al., 2017).

In addition, the smaller variation in B_1_^+^ values across the brain for the London dataset underscores the effect of using different hardware. From this perspective, an important difference between both acquisition sites is the use of single-transmit (Tx) in Maastricht vs. eight channel parallel-transmit (pTx) in London for excitation. Instead of using a single channel RF pulse transmission, pTx makes use of the multiple-transmit coil elements to optimize excitation homogeneity and reduce the B_1_^+^ non-uniformity, particularly important at UHF (Katscher et al., 2003; Zhu, 2004). Although our data show clear differences in the extent of transmit efficiency (i.e., reduced width of B_1_^+^ map histograms), their spatial pattern (i.e., shape) remained relatively comparable: B_1_^+^ is higher in the center of the brain near the limbic lobe, and lower in the temporal lobe, especially. The use of B_1_^+^ shimming using a magnitude least-squares algorithm on the pTx system has the effect (by definition) of reducing the B_1_^+^ variation across the volume of interest and even with a phase-only approach produces a much tighter B_1_^+^ distribution than a single-channel Tx coil. Finally, each acquisition site employs a different gradient coil, corresponding to whole-body vs. head-only, respectively. However, in contrast to pTx, head-only gradients will only have a minor, if not negligible effect on the B_1_^+^ homogeneity. Instead, higher gradient strengths are more useful for cases requiring fast gradient switching such as with functional and diffusion MRI (Uğurbil et al., 2013).

### 4.4. Limitations

The two datasets were acquired using different subjects, and variability in sequence parameters and MRI hardware besides their components determining MP2RAGE B_1_^+^ sensitivity, lowering our precision to pinpoint the B_1_^+^-related biases. However, we were mainly interested in the ‘natural’ variability of T_1_ and segmentation results across independent, though comparable datasets. This more closely matches with typical situations, where datasets are pooled and/or compared. Nonetheless, presence of biological-relevant variability due to differences in population characteristics (e.g., age, sex and handedness distributions) may have attenuated or amplified the observed intra-site differences in cortical T_1_ (Callaghan et al., 2014; Natu et al., 2019) and therefore not necessarily be caused solely by differences in B_1_^+^ sensitivity. Most importantly, the above does not invalidate our conclusions that cortical T_1_ measurements and hippocampal (and subcortical) segmentations can vary substantially due to differences in MP2RAGE acquisition strategy as these statements are based on intra-subject comparisons.

## 5. Conclusions

MRI at 7T holds great promise not only for clinical neuroscientific research but also for assessment of neuroanatomical changes in individual subjects to serve clinical diagnosis. This requires robust acquisition and analysis methods that are insensitive to non-biological variations. In this respect, quantitative imaging approaches such as the MP2RAGE sequence are promising but require extensive validation to optimize their use. Our results show that residual B_1_^+^ effects on MP2RAGE signal intensities, acquired using 7T MRI, not only affect cortical results but impact hippocampal (and subcortical) analyses as well. We confirm that the magnitude of these effects highly depends on the specific set of sequence parameters and/or MRI hardware used. For example, parameters can be chosen to acquire data characterized by improved contrast within cortical or subcortical tissue, or to limit the B_1_^+^ sensitivity of your data. Although the former could be preferred in the case of a specific hypothesis or study aims (i.e., visualization of deep brain nuclei), the latter will greatly improve robustness of the results, promoting comparability across and within subjects and sites. While cortical T_1_, and hippocampal volumes and shape substantially varied between these acquisition strategies initially, it is encouraging that their comparability tends to improve after the post-hoc B_1_^+^ correction. Taken together, our results emphasize the importance of taking into account the presence of potential acquisition-related biases in the data, especially when interpreting small changes in T_1_ or morphology of the brain.

## 6. Acknowledgments

The author Roy AM Haast is supported by BrainsCAN postdoctoral fellowships for this work. Ali R Khan is funded by Canadian Institutes of Health Research Project (366062) and Natural Sciences and Engineering Research Council of Canada Discovery (6639) grants. Infrastructural support was provided by the Canada First Research Excellence Fund to BrainsCAN, Brain Canada, and computational resource through Compute Canada. RF coil technology was supported by a Natural Sciences and Engineering Research Council of Canada Discovery grant (04743) to Ravi S Menon.

## Supplementary Material

**Supplementary Figure 1.**
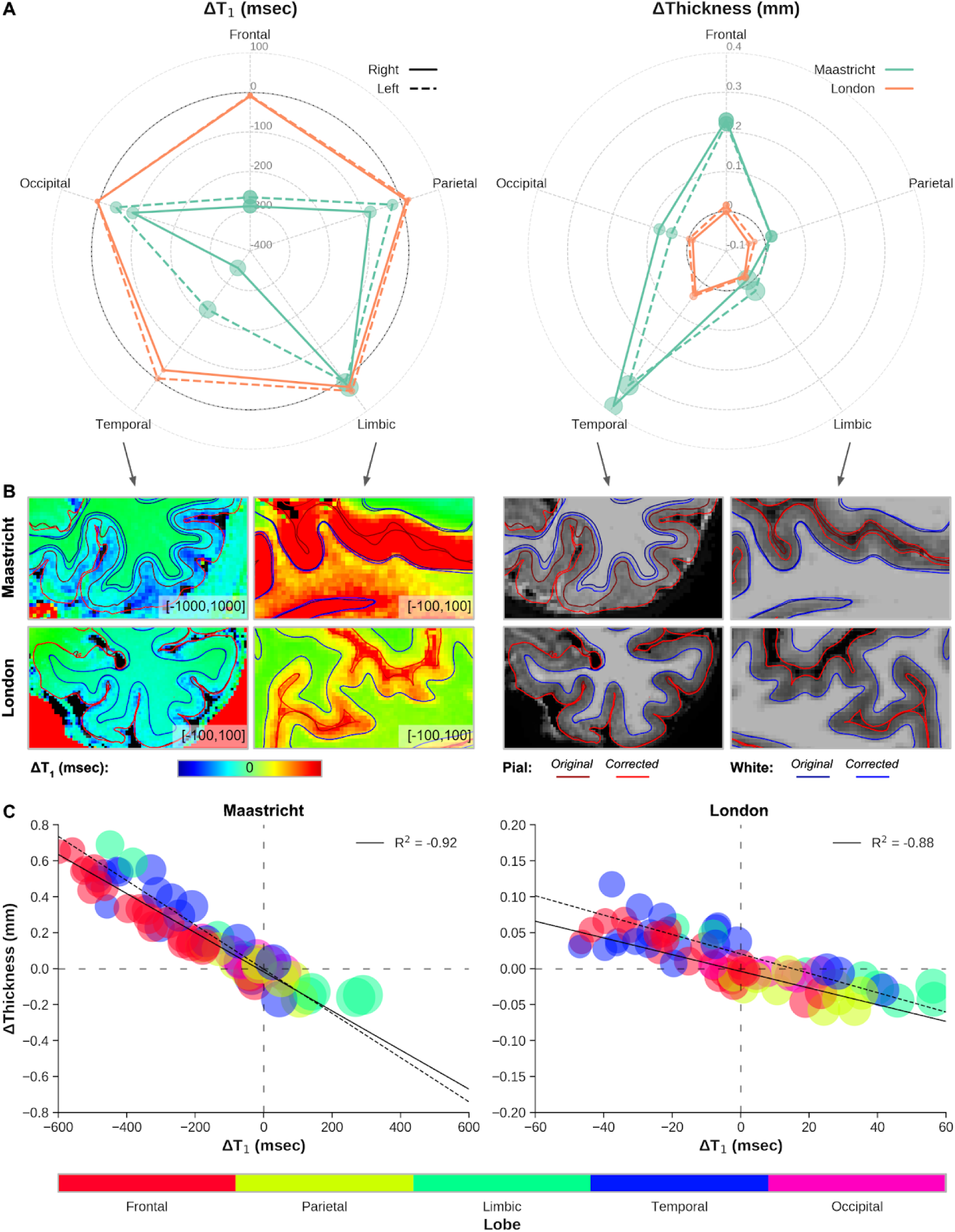
Cortical T_1_ and thickness change as function of cortical lobe. (A) Polar plots showing the average corrected-original T1 (msec, left) and cortical thickness (mm, right) difference for each cortical lobe based on the PALS B12 atlas (i.e., frontal, parietal, limbic, temporal and occipital). Data is shown for both acquisition sites (Maastricht, green and London, orange) and hemispheres (solid vs. dashed). Size of dots represent the (relative) across subjects standard deviation. Dark grey circle indicates x=0 (i.e., no change). (B) Single subject, example Maastricht and London data showing T1 and WM (blue) and pial (red) surface placement before (dark) and after (bright) B1+ correction for temporal and limbic lobes. (C) Change in cortical thickness as function of T1 for each region, color-coded based on cortical lobe, and corresponding correlation coefficients.

**Supplementary Figure 2.**
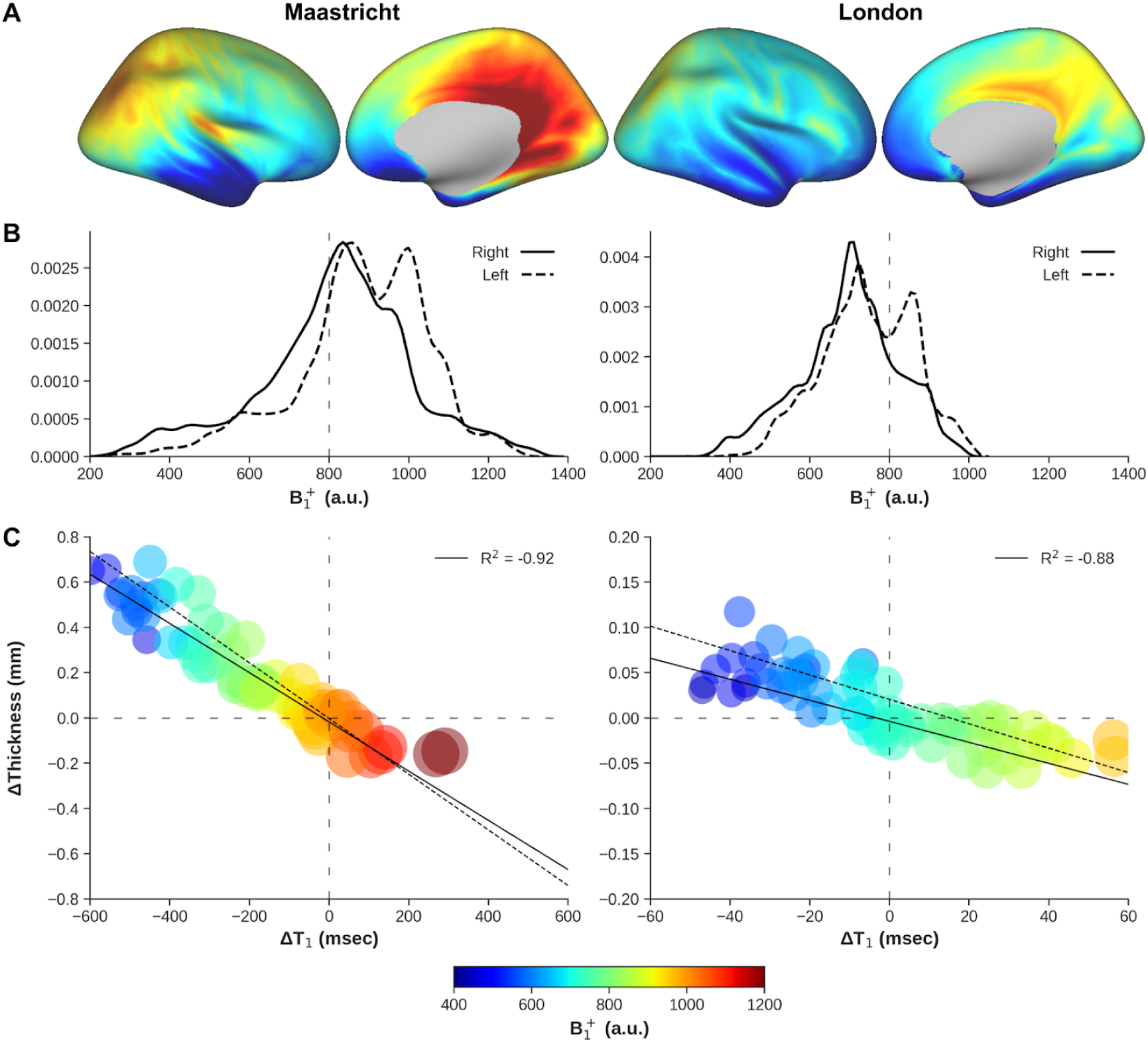
Cortical T_1_ and thickness change as function of cortical B _1_^+^. (A) B_1_^+^ (a.u.) and was mapped onto an inflated right hemisphere surface and averaged across all subjects. (B) Right and left hemispheres (solid and dashed linestyle, respectively) B _1_^+^ distribution. Vertical dashed gray lines are shown for comparison. (C) Change in cortical thickness as function of T_1_ for each region, color-coded based on B_1_^+^, and corresponding correlation coefficients. Left (Maastricht) and right (London) columns represent different acquisition sites. Data is scaled identically for comparison.

**Supplementary Figure 3.**
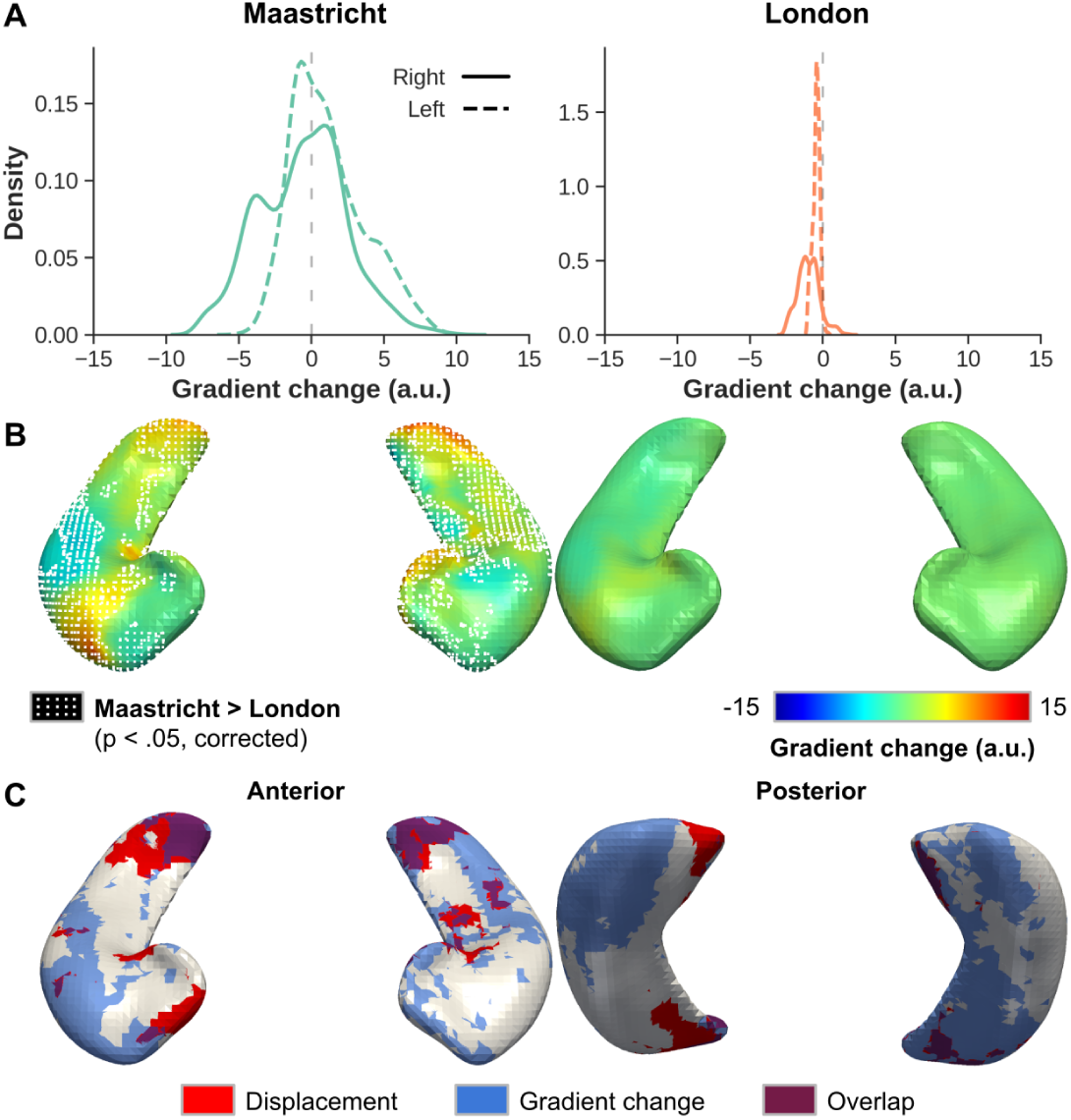
Hippocampal boundary gradient. For each acquisition site (left and right column): (A) Distribution plots showing the average change in gradient magnitude after B_1_^+^ correction at the original hippocampal boundaries. Solid and dashed lines indicate right and left hippocampal data, respectively. (B) Surface representation of average change in gradient for both acquisition sites shown from an anterior perspective. Regions where the Maastricht data was characterized by significantly (p < .05, multiple comparison corrected) larger change in gradient magnitude are demarcated using a dotted pattern. (C) Overlap (in purple) of statistically different vertices based on surface displacement (red) and change in gradient (blue).

**Supplementary Figure 4.**
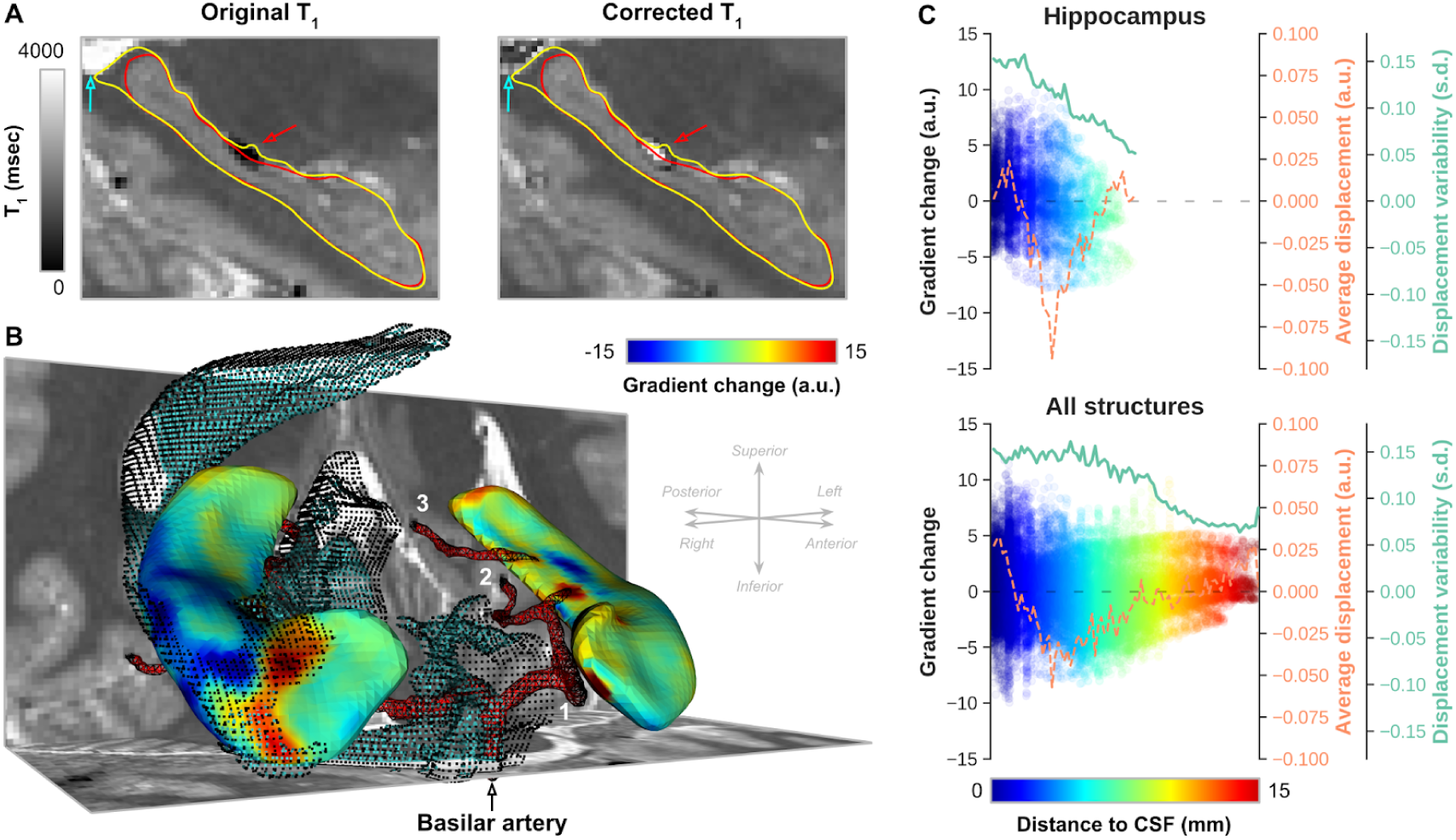
Hippocampal displacement and boundary gradient. (A) Single-subject example from the Maastricht dataset showing the original and corrected T_1_ (msec) maps and corresponding hippocampal segmentation, delineated using yellow and red contours, respectively. Blue and red arrows indicate voxels with large changes in intensity due to the B_1_^+^ correction. (B) For the same subject, 3D scene showing a surface representation of the changes in gradient magnitude along the original hippocampal boundaries (see yellow contours in A), and segmentation of the CSF and major arteries. (C) Plots showing vertex-wise change of gradient magnitude, across subjects average displacement (orange line) and standard deviation (green) as a function of the shortest distance to CSF (color-coded) for hippocampus only (top) and all structures (i.e., including also thalamus, caudate nucleus, putamen, globus pallidum and nucleus accumbens) together (bottom).

**Supplementary Figure 5.**
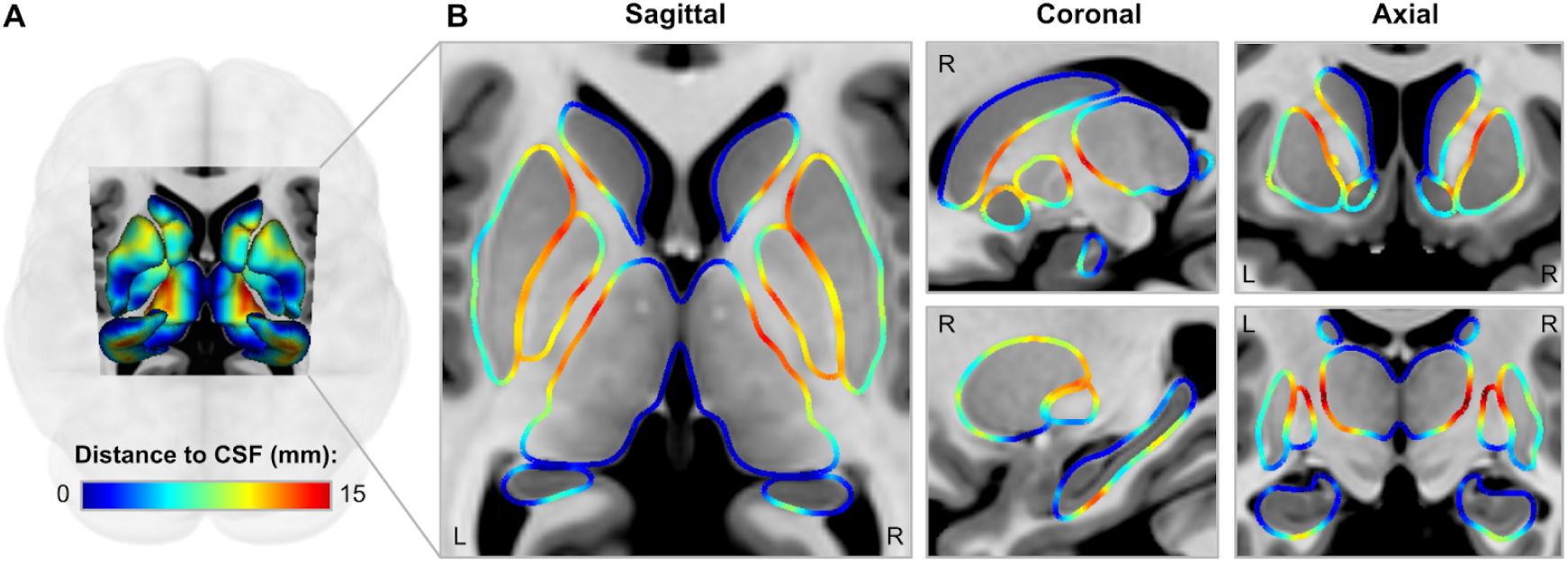
Shortest distance to CSF.

## Supplementary Data 1

### Effect of B_1_^+^ correction on FreeSurfer’s hippocampal subfield analyses

In addition to comparing the standard hippocampal segmentations, we also compared segmentation labels after running the hippocampal subfield analysis pipeline (Iglesias et al., 2015). Supplementary Data Figure 1 allows us to compare between these subfields segmentations generated using the original (left) and corrected (right column) MP2RAGE data for the same single subject within the Maastricht dataset. For illustrative and comparison purposes, we overlaid the surface outlines derived from the hippocampal aseg labels (see also Figure 5A). White arrows in the left column indicate local mismatches between the aseg (i.e., outlines) and subfield (i.e., labels) boundaries, which slightly improve using the B_1_^+^ corrected data (middle column). In general, these mismatches tend to be localized more closely towards the medial and more lateral extents (left and right, respectively, in both the coronal and axial slices) of the CA1 region in the head of the hippocampus, i.e., regions bordered by CSF. Visual comparison of the subfield labels (right column) between original and corrected data demarcate similar locations (along dashed white lines) with varying boundary placement. Finally, compared to those based on the aseg labels (gray dots), a similar trend is observed considering the Dice coefficients (Supplementary Data Figure 1B). However, the Hausdorff distance between boundaries is lower by ± 1.22 voxels.

**Supplementary Data Figure 1.**
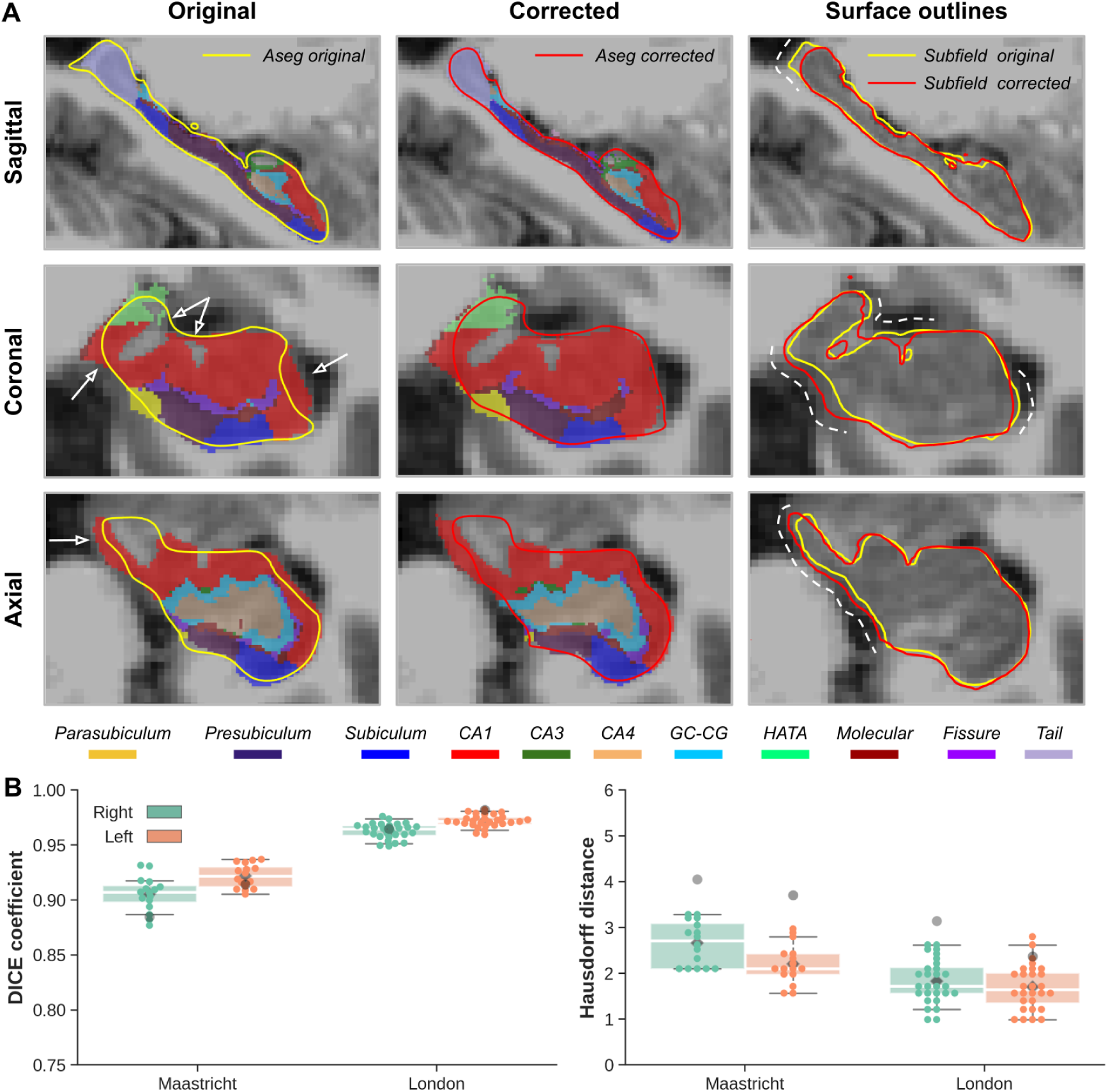
Hippocampal subfields segmentation. (A) Sagittal (top), coronal (middle) and axial (bottom row) cross sections showing a single subject example of left hippocampal subfield segmentation after processing the original (left) and corrected (middle column) MP2RAGE data. Corresponding surface boundaries of the hippocampal aseg segmentations are overlaid for comparison purposes using yellow and red outlines, respectively, while boundaries based on the hippocampal subfield segmentations are shown in the right column. (B) Box plots showing distribution of original vs. corrected hippocampal Dice (left) and Hausdorff (right) scores for both acquisition sites (x-axis), and right (green) and left (orange) hemispheres. Box and whisker extents demarcate interquartile ranges and distribution (excluding outliers), respectively, while diamonds and dots represent group means and individual subjects data, respectively.

## Footnotes

1 Please keep in mind that ‘expected’ refers to the estimated values based on the model presented in Potvin et al. (2016) and our study population characteristics (see also Methods section).

